# Collateral damage of NUDT15 deficiency in cancer provides a cancer pharmacogenetic therapeutic window with thiopurines

**DOI:** 10.1101/2024.04.08.588560

**Authors:** Jacob C. Massey, Joseph Magagnoli, S. Scott Sutton, Phillip J. Buckhaults, Michael D. Wyatt

## Abstract

Genome instability is a hallmark of cancer and are driven by mutations in oncogenes and tumor suppressor genes. Despite successes seen with select targeted therapeutics, this type of personalized medicine is only beneficial for a small subpopulation of cancer patients who have one of a few actionable genetic changes. Most tumors also contain hundreds of passenger mutations that offered no fitness advantage or disadvantage during tumor evolution. Mutations in known pharmacogenetic (PGx) loci for which germline variants encode variability in drug response can cause somatically acquired drug sensitivity. The *NUDT15* gene is a known PGx locus that participates in the rate-limiting metabolism of thiopurines. People with two defective germline alleles of NUDT15 are hypersensitive to the toxic effects of thiopurines. *NUDT15* is located adjacent to the Retinoblastoma (*RB1*) tumor suppressor gene, which often undergoes homozygous deletion in retinoblastomas and other epithelial cancers. We observed that *RB1* undergoes homozygous deletions in 9.4% of prostate adenocarcinomas and 2.5% of ovarian cancers, and in nearly all of these cases *NUDT15* is also lost. Moreover, 44% of prostate adenocarcinomas and over 60% of ovarian cancers have lost one allele of NUDT15, which predicts that a majority of all prostate and ovarian cancers have somatically acquired hypersensitivity to thiopurine treatment. We performed a retrospective analysis of >16,000 patients in the US Veterans Administration health care system and found concurrent xanthine oxidase inhibition (XOi) and thiopurine usage for non-cancer indications is significantly associated with reduced incidence of prostate cancer. The hazard ratio for the development of prostate cancer in patients treated with thiopurines and XOi was 0.562 (0.301-1.051) for the unmatched cohort and 0.389 (0.185-0.819) for the propensity score matched cohort. We experimentally depleted NUDT15 from ovarian and prostate cancer cell lines and observed a dramatic sensitization to thiopurine-induced and DNA damage-dependent toxicity. These results indicate that somatic loss of NUDT15 predicts therapeutic sensitivity to a low cost and well tolerated drug with a broad therapeutic window.

## Introduction

Genome instability is one of the enabling characteristics of cancer^1^ and causes inter- and intra-tumor heterogeneity. Genome instability creates both chaos and opportunities to exploit somatically acquired Achille’s heels that render individual tumors susceptible to existing drugs. Targeted therapeutics such as imatinib and trastuzumab are directed to cancer driver genetic alterations and have notable successes for select patients. Moreover, chaotic genome instability in passenger space sometimes creates neoantigens that enable treatment with immune checkpoint blockade inhibitors. However, despite dramatic successes, resistance often evolves, and immune checkpoint inhibitors do not work for most cancers. Much effort has been expended on therapeutically attacking the consequences of driver mutations; however, cancers contain many more passenger mutations that preceded tumor etiology and are fixed in all clones that may emerge from the first founder driver mutation. In some cases, passenger mutations expand a therapeutic window to existing drugs that could be exploited for cancer therapy.

Pharmacogenetics (PGx) studies have catalogued variability in drug response or toxicity caused by naturally occurring germline variants in drug-metabolizing enzymes. Patient genotype at PGx loci directs clinical therapy and dose selection to achieve therapeutic effect and avoid adverse reactions. We have applied this rationale to cancer genomes, namely, uncovering somatically acquired genetic differences in drug metabolizing enzymes that can be exploited for therapeutic benefit. In this study, we focused on thiopurines and a thiopurine-metabolizing enzyme called NUDT15, for which known germline variants encode clinically relevant increased sensitivity to toxic side effects. The thiopurine 6-mercaptopurine (6-MP) is one of the oldest cancer chemotherapeutics^2^ and it remains a mainstay in leukemia treatments today. Thiopurines such as 6-MP, the prodrug azathioprine, and 6-thioguanine (6-TG) are all metabolically converted to active thioguanine nucleotides (TGNs) and their primary mechanism of action for both therapeutic and toxic effects is via incorporation into DNA as deoxy-6-Thio-GTP. Once incorporated into newly synthesized DNA strands, this mutator analog nucleotide induces apoptosis^3^. NUDT15 (MTH2) is a nucleotide hydrolase (pyrophosphatase) that breaks down nucleoside triphosphates into nucleoside monophosphates^4^. NUDT15 hydrolyzes the ultimate thiopurine metabolite, deoxy-6-TG-TP to prevent 6-TG incorporation into DNA^5^ and thus provides the biochemical basis for the impact of NUDT15 activity on the clinical toxicity of thiopurines^6^. Inactivating germline mutations in *NUDT15* cause patient intolerance to standard doses of thiopurines^7–10^. There are at least 18 germline coding variant alleles of NUDT15 identified to date that compromise NUDT15 biochemical activity and require dose adjustment^11, 12^.

In this study, we determined the extent of NUDT15 loss in cancer genomes and found that, because of the proximity of the *NUDT15* gene to the *retinoblastoma 1* (*RB1*) tumor suppressor gene, many tumors that suffer deep RB1 deletions also lose NUDT15 as collateral damage, including prostate and ovarian cancers. We performed a retrospective analysis of clinical data in the US Veterans Administration VINCI database and determined that patients treated with thiopurines and a xanthine oxidase inhibitor (XOi) for non-cancer indications have a significantly reduced risk of prostate cancer development. The enzyme xanthine oxidase (XO) also inactivates thiopurines^13^; XOi medications such as allopurinol or febuxostat can be used in conjunction with thiopurines to provide clinical effectiveness for low-dose thiopurines. We demonstrate that ovarian and prostate cancer cells in which NUDT15 is experimentally deleted by CRISPR/Cas9 or knocked down by siRNA are sensitized to the toxic effects of 6-TG, and that the mechanism of action requires incorporation into DNA, causes DNA damage and cancer cell death. These results suggest that treating cancer patients who have loss of NUDT15 with thiopurines could be a low cost and well tolerated therapeutic option for thousands of cancer patients.

## Materials and Methods

### Veterans Administration (VA) database analysis

This study utilized data from the US Department of Veterans Affairs (VA). Patient-level medical and demographic information was accessed through the Veterans Affairs Informatics and Computing Infrastructure (VINCI). The study was conducted in compliance with the Department of Veterans Affairs requirements and received Institutional Review Board and Research and Development Approval. Data include veterans with a diagnosis of Crohn’s disease or ulcerative colitis between 2000 and 2022. IBD patients were categorized into three mutually exclusive cohort by their medication usage. Patients who had a 6-MP or azathioprine dispense and a xanthine oxidase inhibitor (XOi) (allopurinol, febuxostat) within 15 days were categorized into the thiopurine+XOi cohort. IBD patients with a thiopurine dispense were included into the thiopurine exposed cohort and those without, categorized into the thiopurine unexposed cohort. Index date was the first date of 6-MP or azathioprine (AZA) dispense. For patients without mercaptopurine or azathioprine the index date was set as the first date of Crohn’s or ulcerative colitis. Patients were followed until first incidence of: 1) prostate cancer, 2) death 3) end of study (31 December 2021). Because treatment assignment was not randomized, a propensity score matching strategy was used to minimize possible selection bias. The propensity score matching model included basic demographic information (age, race, index year) and clinical data (Charlson comorbidity index, BMI, smoking status). We used the propensity score to match thiopurine+XOi patients 1:1 with controls. Matching was conducted sequentially, first by obtaining the matches for the thiopurine exposed cohort and then subsequently for the unexposed cohort. Due to the possibility that mortality could preclude the diagnosis of prostate cancer, a competing risk analysis was conducted. Importantly, systematic differences in mortality between treatment groups could result in biased estimates. Competing risk models evaluate the association between treatment and outcome accounting for the competing risk, in this case, death. Unadjusted analysis includes cumulative incidence estimates generated along with corresponding 95% confidence intervals (CIs). Adjusted Fine and Gray competing risks models are fit, estimating sub-distribution hazard ratios and 95% Cis.

### Cell Culture

OVCAR-8 cells were purchased from the American Type Culture Collection (ATCC) and were maintained in RPMI-1640 and 25 mM HEPES media supplemented with 10% fetal bovine serum and 1% penicillin/streptomycin. HEK 293FT cells were generously provided by Dr. Michael Shtutman and were maintained in DMEM High Glucose media supplemented with 10% fetal bovine serum. 22Rv1 cells were generously provided by Dr. Mengqian Chen and were maintained in RPMI-1640 media supplemented with 10% fetal bovine serum, 1% penicillin/streptomycin, 2mM L-glutamine, 10 mM HEPES, 1 mM sodium pyruvate, 4500 mg/L glucose, and 1500 mg/L sodium bicarbonate. All cells were grown at 37 ℃ in 5% CO_2_. Cells were periodically checked for mycoplasma contamination. 6-Thioguanine was purchased from Sigma-Aldrich (A4882-1G).

### Generation of CRISPR Knockouts and siRNA knockdown treatments

The Cas9 expressing lentiviral vector pCLIP-Cas9-Nuclease-EFS-FLAG-v87 and pCLIP-Dual-SFFV-Zs-Green-Puro clones were from Transomic Technologies. Lentiviral production was carried out as described using pCMV-Δ8.91 and pVSV-G packaging constructs generously provided by Dr. Michael Shtutman.^14^ Lentivirus containing pCLIP-Cas9-FLAG were generated in HEK 293FT cells using PEI as a transfection reagent. Virus-containing media was collected and filtered in a 0.45 μm filter. OVCAR-8 cells were then treated 2x for 24 h each time with virus containing media with 8 μg/mL of polybrene. Cells were the placed under selection with 15 μg/mL of blasticidin (VWR, 10191-144) for 72 h. Confirmation of FLAG-tagged Cas9 expressing cells was confirmed by western blot.

*E. coli* containing the pCLIP-dual gRNA plasmids for NUDT15 (ID: TEDH-1053952) dual-gRNA vectors were acquired from Transomic Technologies via the University of South Carolina Lentiviral Core. The plasmids were isolated using the ZymoPURE II Plasmid Maxiprep Kit (Zymo Research, D4203) according to manufacturer’s instructions. Plasmids were then packaged into lentivirus following the protocol described above. Media containing lentivirus was added to OVCAR-8 cells twice for 24 h each with 8 μg/mL of polybrene. Cells were then placed under selection with 2 μg/mL of puromycin for 96 h. Knockout was confirmed by western blotting and RT-qPCR. The Transomics dual gRNA system allows deletion of the intervening genomic sequence between the two Cas9-mediated DSB sites, which can be detected by PCR as evident by production of a 500 bp product that is only possible if the deletion has occurred between the gRNA B cut site in Exon 1 and the gRNA A cut site in Exon 3.

NUDT15 knock down in 22Rv1 prostate cancer cells was generated using predesigned DsiRNA technology from the IDT TriFECTa® RNAi kit targeting NUDT15 (Integrated DNA Technologies, Design ID hs.Ri.NUDT15.13) Cells were treated with a cocktail consisting of equal parts of three DsiNUDT15 oligonucleotides to a final stock concentration of 1 mM. Cells were treated at a final concentration of 1 nM of the DsiNUDT15 cocktail for 24 h. Cells were then washed once with PBS and allowed to incubate with fresh media for 48 h. Cells were then treated again with 1 nM of the DsiNUDT15 cocktail for 24 h. After the final treatment of DsiNUDT15 cocktail, cells were then trypsinized and plated for further experiments.

### Whole Cell Protein Extraction and Western Blot Analysis

Cells were washed with PBS, trypsinized, collected by centrifugation, and washed once with PBS. The pellet was disrupted in 200 μL of RIPA lysis buffer (VWR, 97063-270) with protease and phosphatase inhibitor cocktail (Thermo Fisher, A32959) and placed on a rocking table at 4 ℃ for 30 min. The samples were sonicated three times (10 seconds on/5 seconds off) at 40% amplitude and rested on ice for 5-10 minutes. Protein concentrations were determined by BCA assay (Thermo Fisher, 23225). Protein samples 30 μg) were separated by SDS-PAGE and transferred to a PVDF membrane overnight. All membranes were blocked with 5% milk blocking buffer (Bio-Rad) in PBST (1x PBS with 0.05% Tween-20) or TBST (1x TBS with 0.05% Tween-20) prior to primary antibody addition, which after 1 h blocking occurred overnight at 4 ℃ on a plate rocker. The following primary antibodies were used: phospho-γH2AX (Abcam, ab11174, 1:5000), NUDT15 (ABclonal, A8368, 1:2000), GAPDH (Cell Signaling, 5174S, 1:2000), β–Tubulin (Novus Biologicals, NB600-936, 1:2000). After overnight incubation, membranes were washed with PBST or TBST three times for ten min. Anti-rabbit-HRP secondary antibody (Sigma Aldrich, NA9340V, 1:2000) was added in 5% milk and allowed to incubate at RT for 2 h on a plate rocker. The membranes were washed with PBST or TBST three times for ten min. After addition of ECL blotting substrate (Pierce, 32106), membranes were imaged using the ChemiDoc Touch Imaging System (Bio-Rad). Quantification of bands used Bio-Rad Image Lab software normalized to β–Tubulin expression.

### Metaphase Spreads

Cells were plated at a density of 30,000 cells per well and allowed to incubate at 37 ℃ overnight. Cells were treated for 6 h, followed by a PBS wash and addition of 2 mL fresh media, and incubation for 18 h at 37 ℃. After 18 h, media containing 20 ng/mL of colcemid was added and further incubated for 4 h. After incubation, cells were trypsinized and placed in a hypotonic solution (2 mg/mL KCl) for 10 min at RT. Cells were pelleted and resuspended twice in an ice-cold fixative solution containing a 3:1 mixture of methanol to glacial acetic acid. To prepare chromosome spreads, cells were dropped by pipet onto slides, air dried, and stained in 10% Giemsa (Sigma), washed with water and allowed to air dry overnight. Slides were scored blinded on a Nikon Eclipse E600 Microscope using the 100x objective under oil. Chromosomal aberrations were scored with a minimum of 950 chromosomes per sample.

### Colony Forming Assay

Cells were plated at a density of 1000 cells per well in a 6 well plate 24 h prior to drug treatment. After treatments, cells were incubated for 9 days, then fixed in 100% methanol for 10 min. After methanol removal and plates were air dried, the colonies were stained with filtered 10% Giemsa stain and imaged with the ChemiDoc Touch Imaging System (Bio-Rad). The area of colonies in each well were quantified using the Colony Area Plugin for ImageJ.^15, 16^

### Competitive Growth Assays

Cells were seeded in 6-well plates at a density of 80,000 cells per well with a 50-50 mixture of wild type and NUDT15 knockout cells and treated with LD_50_ and LD_10_ doses of 6-TG. The cells incubated in a BioTek BioSpa 8 live-cell imager at 37 ℃ with 5% CO_2_ for 72 h with images at 4x magnification taken every 6 h until the next passage or end of treatment. Images captured were phase contrast and GFP fluorescent filter images. After 72 h, cells were split 1:5 and added to a new plate containing fresh media. Treatments and visualization were repeated for a total of five passages. The images were compiled with GFP^+^ versus the total number of cells quantified over time.

### Flow Cytometry

Cells were collected and washed with 2 mL of ice cold 1x PBS with 1% FBS and cells were pelleted at 1100 *x g* for 5 min. The supernatant was removed, and cells were then gently resuspended in 2 mL of ice cold 1x PBS. Two milliliters of ice cold 100% ethanol were added dropwise, then 3 mL of ice cold 100% ethanol was added to a final volume of ∼70%. The cells were then stored at −20 ℃ for at least one night.

To determine the cell cycle profile, cells were pelleted, the supernatant was removed, and cells were washed with 2 mL of 1x PBS + 1% FBS. The cells were centrifuged, the supernatant was removed, and cells were then gently resuspended in 1 mL of 1x PBS with 3 μM of DAPI and allowed to incubate for 15 minutes at room temperature in the dark. The samples were run on a BD LSR II Flow Cytometer in the Microscopy and Flow Cytometry Facility at the University of South Carolina, College of Pharmacy. At least 10,000 cells were analyzed per experiment.

### Alkaline Comet Assay

Cells were plated in a 24 well plate at a density of 1000 cells per well and allowed to incubate for 24 h. Cells were then treated over the course of 72 h. After treatment, cells were trypsinized, washed with PBS, and stored on ice. Cells were collected in 30 μL of PBS at a density of 1 x 10^5^ cells/mL and mixed with 300 μL of molten 1% low melting agarose in PBS at 37 ℃. Using pre-coated slides that were warmed to 37 ℃ (Trevigen, 4252-040-01), 50 μL of the cell and agarose mixture was added and spread evenly with the side of the pipet tip. Slides were then stored at 4 ℃ in the dark for 20 min. Slides were then immersed in the dark in pre-chilled lysis buffer (Trevigen, 4250-010-01) for at least 1 h to no longer than overnight. The lysis buffer was then removed and replaced with freshly made, pre-chilled unwinding solution (200 mM NaOH, 1 mM EDTA, pH > 13) for 1 h at 4 ℃ in the dark. Slides were then moved into the Trevigen CometAssay ES unit with freshly made, pre-chilled alkaline running solution (200 mM NaOH, 1 mM EDTA, pH > 13) and run at 21 V for 20 min according to manufacturer’s recommendations. The slides were then immersed in RT ddH_2_O twice for 5 min each and RT 70% EtOH for 5 min. The slides were then allowed to completely dry overnight at 37 ℃ before imaging.

For imaging, slides were stained with 1x SYBR Gold in 1x TE buffer for 30 minutes and de-stained with ddH_2_O. The cells were allowed to dry for 20 min at 37 ℃ in the dark. The cells were imaged using the Carl Zeiss LSM Confocal Microscope camera using Zen Pro image processing system at a 10x magnification. The images were then scored using Trevigen Comet Analysis Software. At least 50 comets were scored per treatment group. Statistical significance was determined using two-way ANOVA with GraphPad Prism.

## Results

### NUDT15 loss is collateral damage in RB1-deleted cancers

Screening for NUDT15 variants with deficient nucleotide hydrolase activity is necessary prior to starting a patient on thiopurine treatments, so that dosing adjustments can minimize patient toxicity and maximize therapeutic benefit^17^. We asked whether cancer genomes contain deficiencies in *NUDT15*, which would provide a therapeutic window for patients with wild-type NUDT15 activity. The *NUDT15* gene on chromosome 13 is within 300 kb from the *retinoblastoma 1* (*RB1*) tumor suppressor gene (*NUDT15:* 48,037,726 - 48,052,755, *RB1*: 48,303,751 - 48,481,890). Evidence found through publicly available databases shows that *NUDT15* copy number is linked to changes in copy number of the cell cycle regulating protein *RB1* across many tumor types, with prostate adenocarcinoma showing the highest frequency (**Figure 1A**). TCGA copy number data (GISTIC) shows that almost half of prostate cancers have copy number losses to NUDT15, of which 9.4% are deep deletions and all but one of which are co-deleted for *RB1* (**Figure 1B**, left panel). ^18, 19^ There is a strong positive correlation between *RB1* and *NUDT15* gene deletion in the 489 prostate adenocarcinoma samples examined (p-value <0.001, see methods). It therefore appears that tumors suffering deep genomic deletions at the *RB1* tumor suppressor also lose *NUDT15*. In prostate cancers, deletion of *NUDT15* is strongly associated with loss of NUDT15 expression (**Figure 1C**, left panel). Detailed analysis of the ovarian serous cystadenocarcinomas in TCGA revealed almost identical findings. TCGA copy number data (GISTIC) shows that over 60% of ovarian cancers have copy number losses to NUDT15, all but one of which are co-deleted for *RB1* (**Figure 1B**, right panel). There is a strong positive correlation between *RB1* and *NUDT15* gene deletion in the 539 ovarian cancer samples examined (p-value <0.001, see methods). The expression data showed deletion of *NUDT15* strongly associated with loss of NUDT15 expression in tumors with both shallow and deep deletions (**Figure 1C**, right panel). Collectively, the tumors that lost NUDT15 expression because of genomic deletion are therefore permanently defective in NUDT15 enzyme activity, as reversion mutations from homozygous deletions are extremely unlikely.

**Figure 1.**
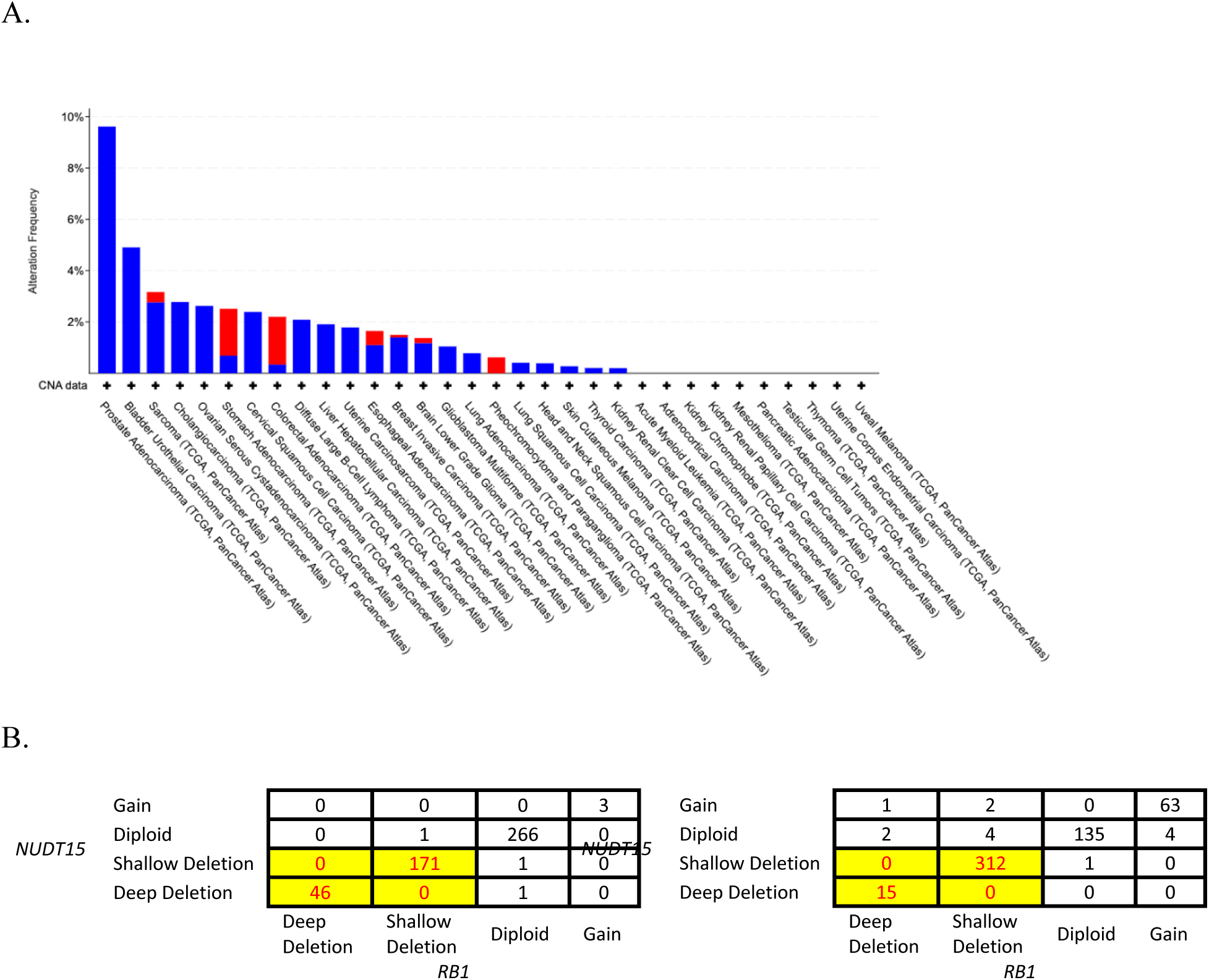
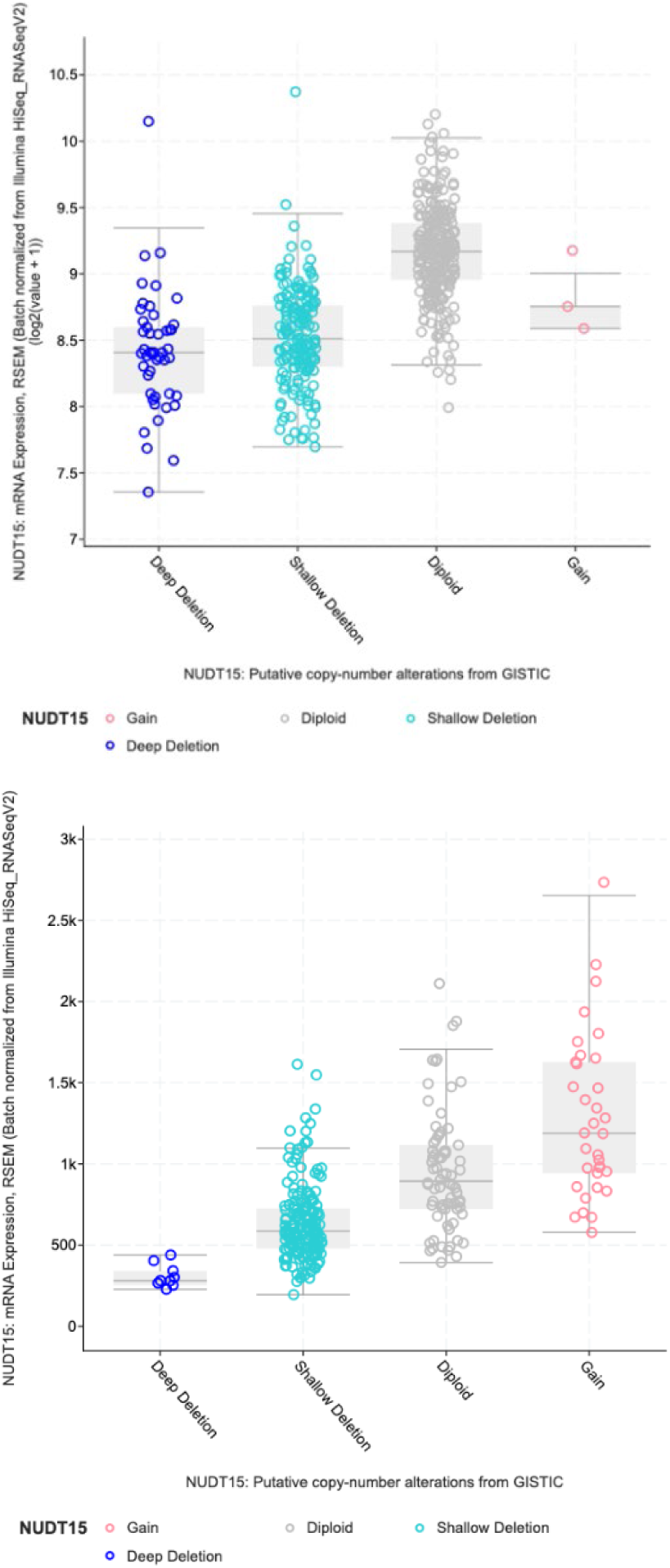
Association of NUDT15 and RB1 alterations in human cancers. **A.** Analysis of *NUDT15* Deep Deletions in the TCGA. Blue represents deep deletion events. Red represents amplification events. **B.** Co-occurrence of *NUDT15* and *RB1* deletions in prostate cancers (Left panel) and ovarian (Right panel) **C.** loss of *NUDT15* expression (y-axis) in cancers with *NUDT15* copy number loss. Left panel, prostate cancer. Right panel, ovarian cancer. Dark Blue represents cancers with deep deletion events. Cyan represents cancers with shallow deletion events. Grey represents cancers with diploid NUDT15 copy number. Red represents cancers with *NUDT15* copy number gains.

### Thiopurines prescribed for ulcerative colitis or Crohn’s disease are associated with a reduced risk of prostate cancer

Thiopurines are used as immunosuppressants in the management of ulcerative colitis and Crohn’s disease. The enzyme xanthine oxidase (XO) also inactivates thiopurines^13^. We analyzed the incidence of prostate cancers in a VA cohort of Crohn’s disease patients that were unexposed to thiopurines or exposed to the combination of thiopurines and the xanthine oxidase inhibitors (XOi) allopurinol or febuxostat, which are used to manage symptoms of gout. Table 1 displays the baseline patient characteristics for the unadjusted cohort and a subset cohort made of patients matched by propensity scores. After propensity score matching, each cohort had 416 patients and baseline patient characteristics were more closely matched. Patients treated with both thiopurines and an XOi had significantly decreased incidence of prostate cancers (**Table 1**). Estimating Fine and Gray competing risks models revealed that among the propensity score matched cohorts, patients treated with both thiopurines and an XOi had a significantly lower hazard of prostate cancer compared to patients unexposed to thiopurines (**Table 2**). The hazard ratio for the development of prostate cancer in patients treated with both thiopurines and xanthine oxidase inhibitors was 0.562 (0.301-1.051) for the unadjusted cohort and 0.389 (0.185-0.819) for the propensity score matched cohort.

**TABLE 1.** Baseline patient characteristics for the unadjusted cohort and a subset cohort made of patients matched by propensity scores.

|  |  | Unadjusted Cohort |  |  |  | Cohort of Propensity-score matched people |  |  |  |
| --- | --- | --- | --- | --- | --- | --- | --- | --- | --- |
| Variable |  | Thiopurine<br>unexposed<br>N=107100 | Thiopurine<br>exposed<br>N=12760 | Thiopurine+<br>XOi N=416 | P.<br>value | Thiopurine<br>unexposed<br>N=416 | Thiopurine<br>exposed<br>N=416 | Thiopurin<br>e+ XOi<br>N=416 | P.value |
| Age |  | 60.4<br>(15.58) | 54.34 (16.27) | 60.7 (12.75) | <0.001 | 61.01<br>(13.09) | 60.53 (13.51) | 60.7<br>(12.75) | 0.866 |
| Race | Black | 11949<br>(11.16%) | 1210 (9.48%) | 32 (7.69%) | <0.001 | 21 (5.05%) | 21 (5.05%) | 32<br>(7.695%) | 0.024 |
|  | Other/<br>unknown | 12326<br>(11.51%) | 1239 (9.7%) | 39 (9.35%) |  | 44<br>(10.58%) | 23 (5.53%) | 39<br>(9.38%) |  |
|  | White | 82825<br>(77.33%) | 10311(80.81%) | 345<br>(82.93%) |  | 351<br>(84.38%) | 372 (89.42%) | 345<br>(82.93%) |  |
| Charlson comorbidity<br>index |  | 1.04 (1.93) | 0.68 (1.28) | 1.24 (1.77) | <0.001 | 1.22 (1.89) | 1.15(1.83) | 1.24(1.77) | 0.758 |
| BMI | <18.5 | 1586<br>(1.48%) | 258 (2.02%) | <5(#) | <0.001 | <5(#) | <5(#) | <5(#) | 0.476 |
|  | 18.5-24.9 | 24559<br>(22.93%) | 3539 (27.74%) | 56<br>(13.46%) |  | 62<br>(14.90%) | 42 (10.1%) | 56<br>(13.46%) |  |
|  | 25-29.9 | 40489<br>(37.81%) | 4996 (39.16%) | 140<br>(33.65%) |  | 142<br>(34.14%) | 145 (34.86%) | 140<br>(33.65%) |  |
|  | 30+ | 39352<br>(36.74%) | 3912(30.66%) | 215<br>(51.68%) |  | 210<br>(50.48%) | 225 (54.09%) | 215<br>(51.68%) |  |
|  | Missing | 1114<br>(1.04%) | 55 (0.43%) | <5(#) |  | <5(#) | <5(#) | <5(#) |  |
| Smoker |  | 8379<br>(7.82%) | 1100 (8.62%) | 22(5.29%) | <0.001 | 22 (5.29%) | 19 (4.57%) | 22<br>(5.29%) | 0.86 |
| Index year |  | 2011.84<br>(6.88) | 2009.74 (6.14) | 2012.25<br>(5.91) | <0.001 | 2012.3<br>(6.05) | 2011.95<br>(5.75) | 2012.25<br>(5.91) | 0.659 |

**TABLE 2.** Cumulative incidence of prostate cancer: estimates and 95% CI.

| Time window | Unadjusted Cohort |  |  |  | Propensity score matched Cohort |  |  |  |
| --- | --- | --- | --- | --- | --- | --- | --- | --- |
|  | Thiopurine unexposed | Thiopurine exposed | Thiopurine+ XO<sub>i</sub> | Gray's test P.value | Thiopurine unexposed | Thiopurine exposed | Thiopurine+ XO<sub>i</sub> | Gray's test P.value |
| 10 year risk | 5.0% (4.8%,5.1%) | 4.1% (3.7%,4.5%) | 3.0% (1.5%,5.3%) | 0.004 | 6.8% (4.3%,10%) | 3.8% (2.1%,6.2%) | 3.0% (1.5%,5.3%) | 0.005 |
| 20 year risk | 7.2% (7.0%,7.4%) | 6.9% (6.3%,7.5%) | 3.0% (1.5%,5.3%) |  | 8.5% (5.3%,13%) | 10% (4.9%,18%) | 3.0% (1.5%,5.3%) |  |

### NUDT15 depleted cells are sensitized to a thiopurine therapeutic

Because of the striking findings regarding NUDT15 loss discussed previously, we sought to confirm whether direct loss of NUDT15 in ovarian and prostate cancer cells would cause sensitization to a model thiopurine and whether the mechanism of action would be consistent with that observed for thiopurine treatments. Successful deletion of the *NUDT15* gene was confirmed in the ovarian OVCAR-8 cells by PCR amplification of a 500 bp product that is only possible if the deletion occurred (**Figure 2A**, left panel). Western blot analysis of the parental and NUDT15 KO cells showed near complete loss of the NUDT15 protein (**Figure 2b**, right panel). Parental control and NUDT15 KO cells were treated with 6-TG to determine the extent of NUDT15-dependent sensitization to thiopurines in the ovarian cancer cell line. 6-TG was chosen because it is most directly metabolized to the ultimate d-6-TGTP metabolite through which all thiopurines are metabolized. The NUDT15 KO cells were >15-fold more sensitive to 6-TG treatments with an LC_50_ of 0.28 μM compared to an LC_50_ of 4.32 μM for the NUDT15 proficient cells (Figure 3A). Increased sensitivity to 6-TG was also seen in 22Rv1 prostate cancer cells with NUDT15 expression depleted via NUDT15 siRNA. The NUDT15-deficient prostate cancer cells were approximately twice as sensitive to 6-TG with an LC_50_ of 1.48 μM compared to 3.13 μM in the scrambled siRNA control (**Figure 3B**).

**Figure 2.**
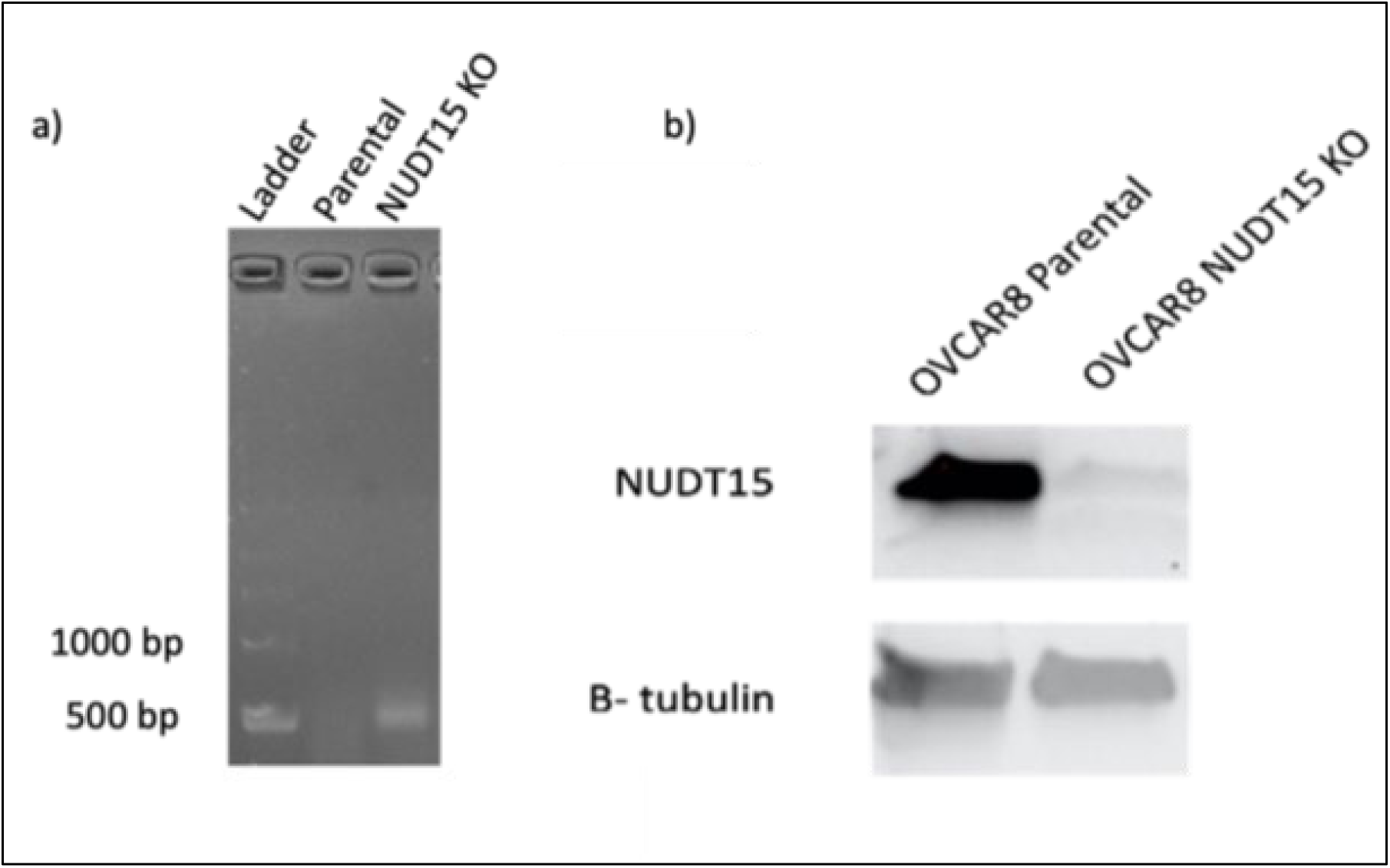
Confirmation of NUDT15 CRISPR Knockout in Ovarian Cancer Cells. **A)** PCR Amplification of a 500 bp product from isolated genomic DNA with primers that flank the location of the dual-sgRNA target sequences, confirming genomic knockout of the NUDT15 gene. The The left lane shows the 1 kb ladder. The middle lane shows a sample of genomic DNA subject to PCR conditions and no discernable PCR product detected. The right lane shows the presence of the 500 bp product formed in the NUDT15-deficient cell line. **B)** Western blot analysis of whole cell lysates from parental and NUDT15 knockout cells. β-tubulin served as the loading control. Representative images three experiments shown.

**Figure 3.**
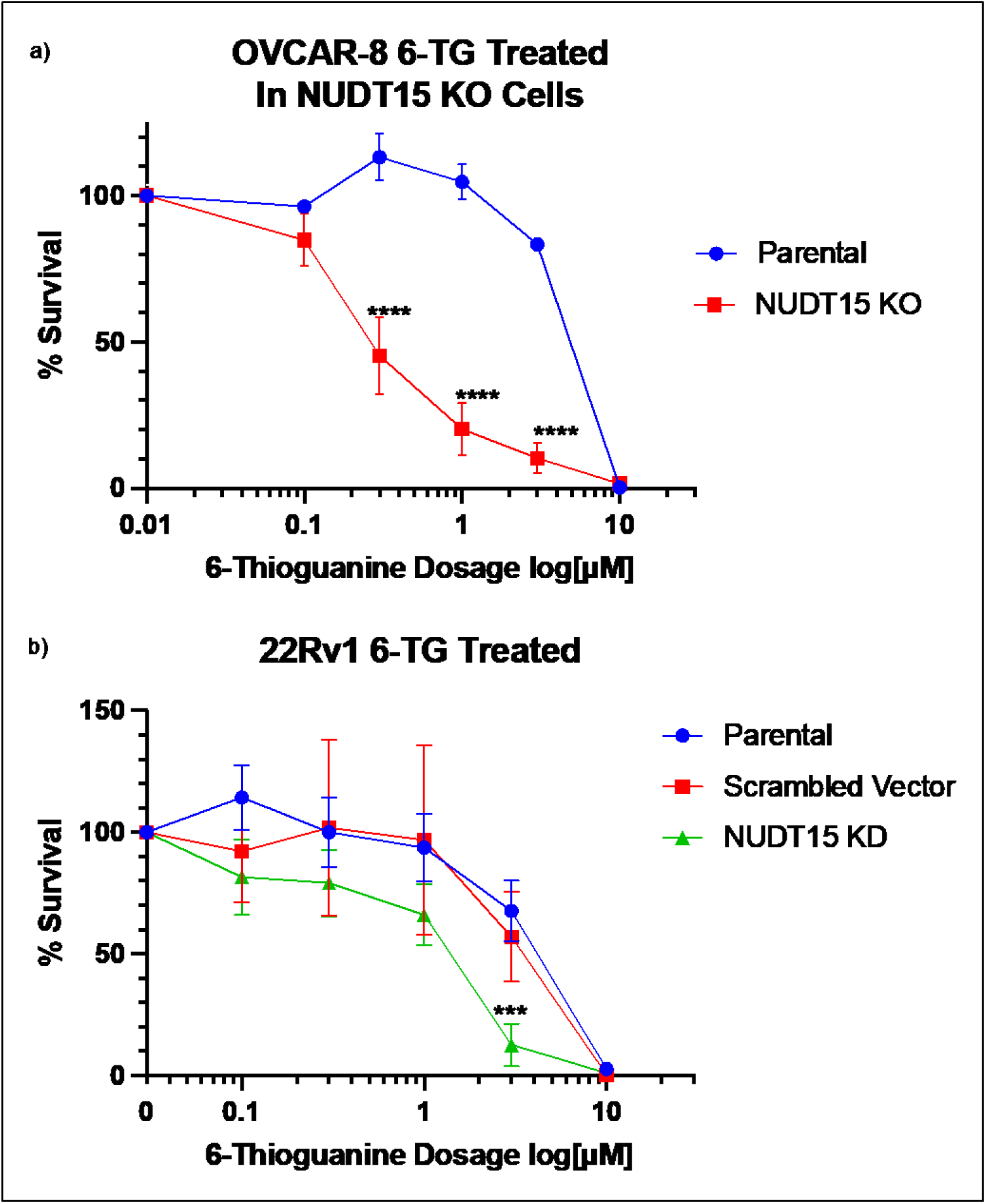
NUDT15 KO and KD cells show sensitivity to 6-thioguanine treatment. Colony forming assays of cells treated with 6-TG at the indicated doses. **A)** OVCAR-8 CRISPR KO cells and **B)** 22Rv1 prostate cancer cells transiently transfected with siRNA against NUDT15. Survival is plotted compared to untreated control. Each data point is a mean of 3 independent experiments. Statistical significance was determined by two-way ANOVA *** p< 0.001 **** p < 0.0001

Selection assays were conducted using live-cell imaging, in which NUDT15-proficient and deficient OCVAR-8 cells were co-incubated and treated with 6-TG (**Figure 4**). Because the gRNA plasmid also contains a GFP cassette, the NUDT15-deficient cells are GFP^+^, which provides a marker to track the fate of NUDT15-deficient cells relative to NUDT15-proficient cells during 6-TG treatments. The percentage of GFP^+^ cell area in the non-treated cells (blue line) at the start of each passage was ∼0.2, and by passage four and five the GFP^+^ cell area increased to 0.4, showing that NUDT15-deficient OVCAR-8 cells do not suffer from any growth defects relative to NUDT15-proficient cells. Cells were treated with successive rounds with 0.1 μM and 0.3 μM 6-TG for 72 h, representing LC_10_ and LC_50_ doses in the NUDT15-deficient cells, for a total of 5 passages. **Figure 4A** shows representative phase contrast and green-fluorescent images of the cells after passages 1, 3, and 5. **Figure 4B** shows the ratio of GFP^+^ cells across the 72 h time course for each of the five passages. After the first two passages, the percentage of GFP^+^ cell area was similar (**Figure 4A, 4B**). Starting with Passage 3, the ratio of GFP^+^ cell area representing NUDT15-deficient cells decreases in both 0.1 and 0.3 μM 6-TG-treated populations (**Figure 4B**, Top Right). The loss of GFP^+^ cell area continued with Passage 4 in 6-TG treated cells (p < 0.01) (Figure 4B, Bottom Left). At Passage 5, GFP^+^ cell area representing NUDT15 KO are almost completely abolished after treatments with 6-TG (**Figure 4A**, Bottom Row, **4B**, bottom right, both p < 0.0001). A slight loss of overall viability was noted in Passage 5 cells treated with 0.3 μM of 6-TG (compare NT and 0.1 to 0.3 mM in the bottom right of **Figure 4A**), showing that the higher dose of 6-TG used here was mildly anti-proliferative to the NUDT15-proficient cells.

**Figure 4.**
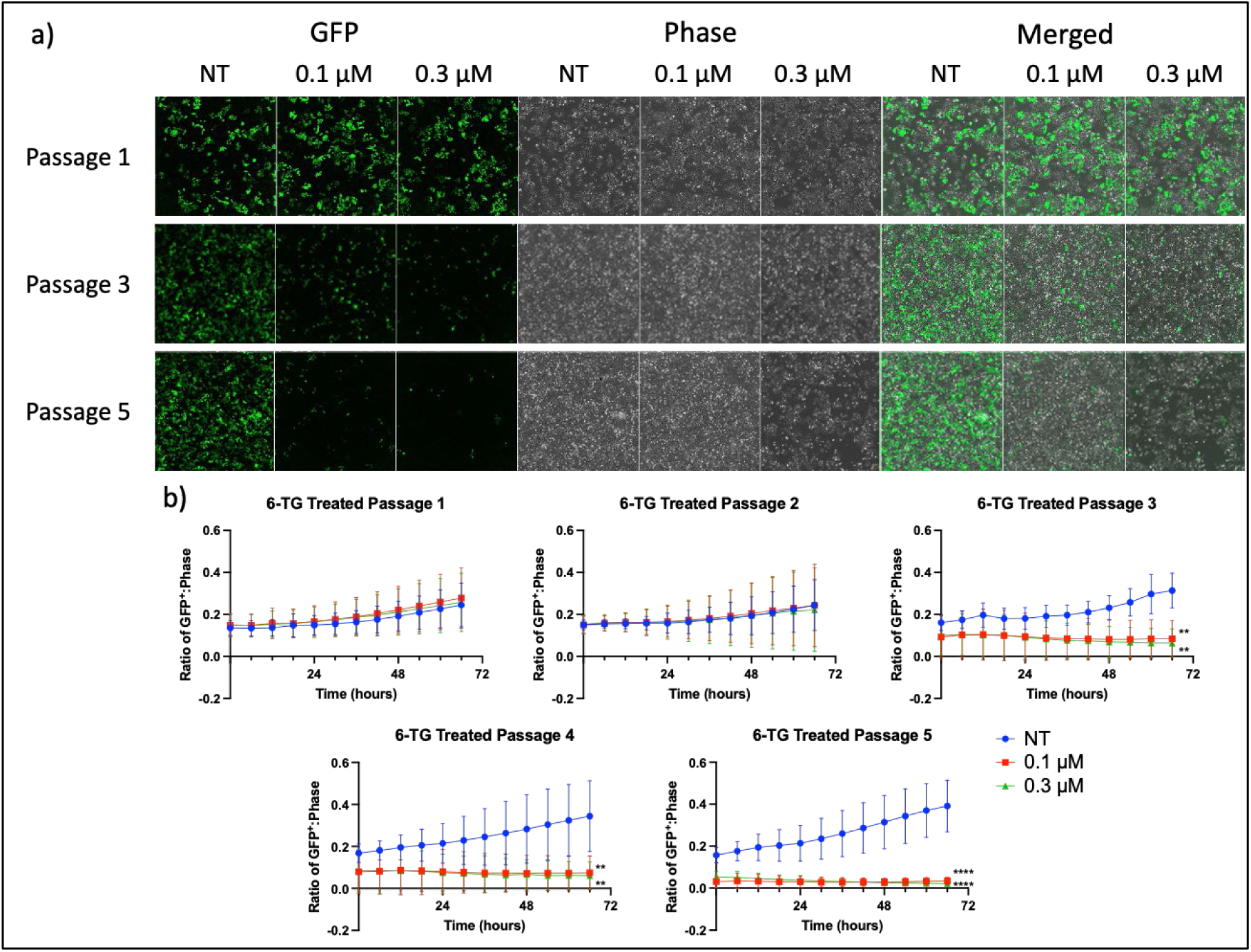
Competitive growth assay of NUDT15 proficient and GFP^+^ NUDT15 KO cells after 6-TG treatment. Cells were imaged every 6 h during the course of a 66 h exposure to 6-TG prior to re-plating for fresh drug exposure in subsequent passages. **A)** Representative images of Passage 1, Passage 3, and Passage 5 cells were taken after 66 hours of treatment. Images show GFP (left), phase contrast (middle), and the merged image (right). **B)** Quantification of the ratio of GFP^+^ cells at each time point relative total cell number calculated by phase contrast with each group. Significance was determined by two-way ANOVA. ** p < 0.01 **** p < 0.0001

### NUDT15-depleted cells suffer cell cycle arrest and markers of DNA damage

Previous studies have shown that 6-TG induces a G_2_/M phase arrest in cells mediated by MMR ^20–22^. To determine whether NUDT15 deficiency heightens the 6-TG mediated G_2_/M phase arrest, cells were treated, and cell cycle distribution was monitored at 24, 48, and 72 h. No significant difference between the parental cells and the NUDT15 KO cells was observed after 24 h treatment (data not shown), in agreement with prior studies showing that one round of replication is required to incorporate 6-TG into the genome. After 48 h 6-TG treatment, an almost 2-fold increase in the S-phase population was observed in NUDT15 KO cells treated with 0.3 μM 6-TG, compared to untreated (NT) NUDT15 KO cells (**Figure 5A**). The change in S phase cell distribution was also significantly different (p < 0.001) compared to the NUDT15-proficient cells at the same concentration (**Figure 5A**, tan bars 1-3). After 72 h 6-TG treatment, the two-fold increase in S-phase NUDT15 KO cells at the 0.3 μM dose persisted (**Figure 5B**, tan bars 4-5). A two-fold increase in accumulation of G2/M cells was observed at the 1.0 μM dose in the NUDT15 KO cells compared to the NUDT15-proficient cells (**Figure 5B**, blue bars 4-6). This supports previous findings that cells progress though mitosis with doses of 6-TG, while higher doses of 6-TG cause a G_2_ arrest.^23^

**Figure 5.**
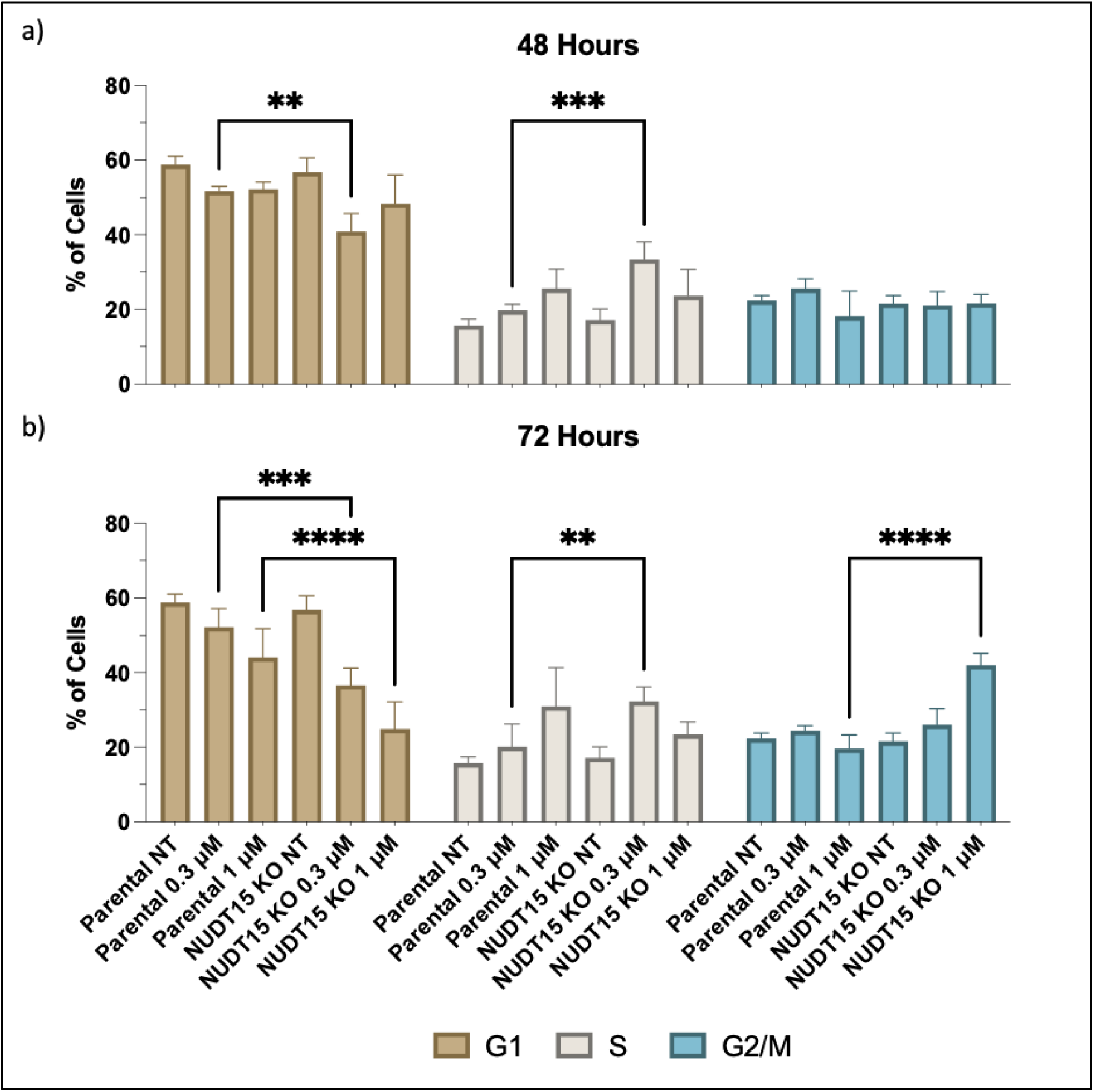
Cell-cycle profile of parental and NUDT15-deficient cells in response to 6-TG treatments. Flow cytometry results of cells treated with 6-TG at the indicated doses for the indicated time. Each panel shows the G_1_, S phase, G_2_/M phase populations of parental and NUDT15 KO cells after exposure to 6-TG for: **A)** 48 h and **B)** 72 h. Each bar represents three independent experiments. Statistical significance was determined by two-way ANOVA. ** p < 0.01, *** p < 0.001, **** p < 0.0001

Induction of the DNA damage marker γ-H2AX was assessed by western blotting after 6-TG treatments in parental and NUDT15-deficient OVCAR-8 cells. Following treatments of 0.3 μM and 1 μM of 6-TG, increases in γ-H2AX are induced in the NUDT15 KO cells after 48 and 72 h exposure to 6-TG (**Figure 6A**, lanes 7 and 8). Quantitation shows there is a two-fold increase in γ-H2AX in the NUDT15 KO cells (red bars) compared to the parental cells (blue bars) following 0.3 μM 6-TG treatments at 48 and 72 h (**Figure 6B**). NUDT15 KO cells treated with 1 μM 6-TG showed 4.5 and 4-fold increases at 48 h and 72 h, respectively, which were substantially higher compared to parental cells (**Figure 6C**).

**Figure 6.**
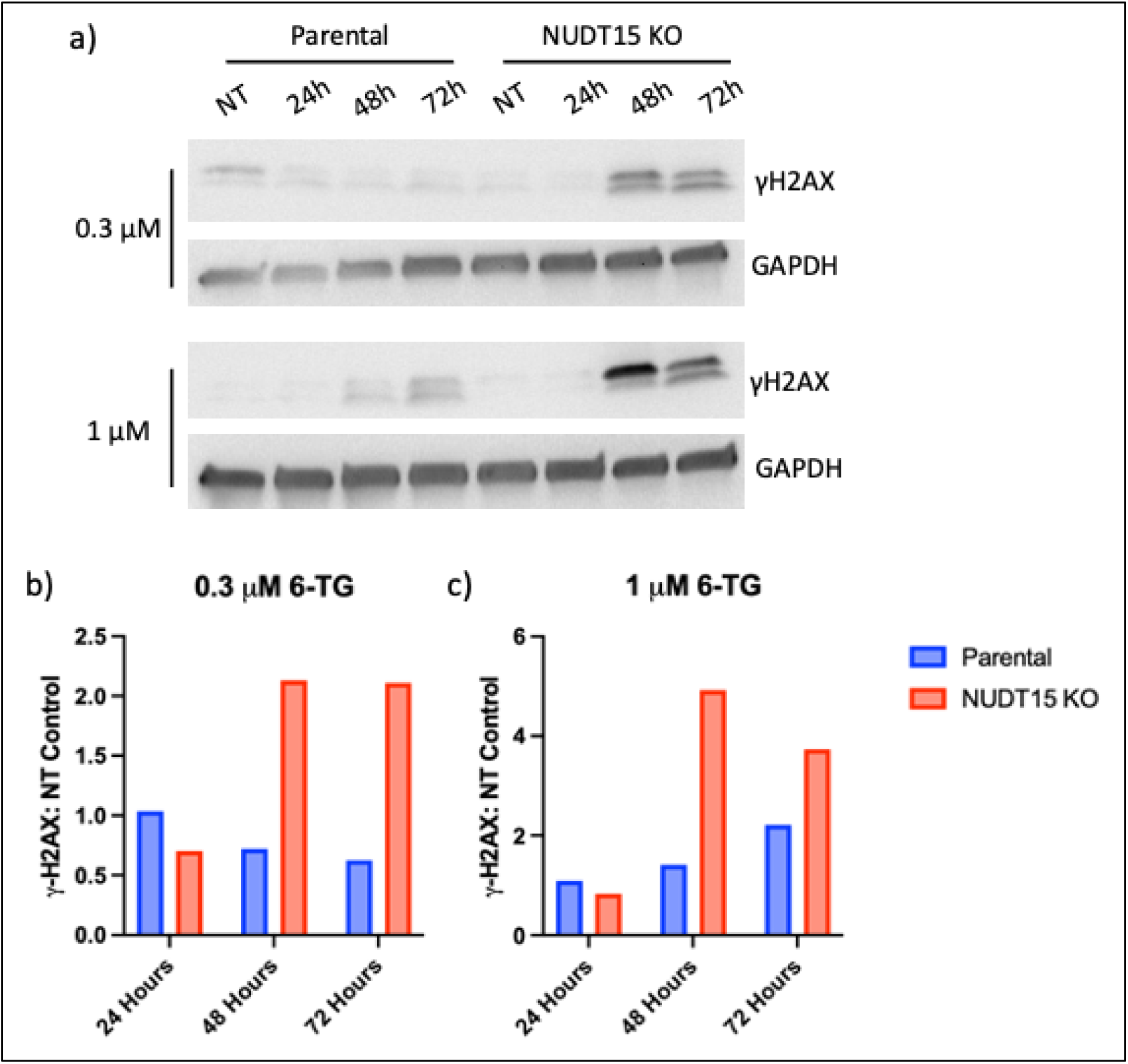
Induction of DNA Damage marker γ-H2AX after 6-TG treatment. Western blots measuring γ-H2AX in NUDT15 parental and KO cells after 6-TG treatment. **A)** The γ-H2AX signal after 6-TG treatment of 0.3 μM and 1 μM doses for 24, 48, and 72 h in parental cells (lanes 1-4) and NUDT15 KO cells (lanes 5-8). **B-C**) Quantification of γ-H2AX bands normalized to GAPDH after 0.3 μM **(B)** or 1 μM **(C)** of 6-TG. Bars show total band volume of each lane compared to NT control cells.

Alkaline comet assays were performed to quantitate DNA single and double strand breaks following 6-TG treatment (**Figure 7**). NUDT15-proficient (blue bars) and deficient (red bars) cells were treated with a 0.3 μM or 1 μM dose of 6-TG for 24, 48, and 72 h and analyze for percentage of DNA in the tail. At 24 h, the NUDT15-proficient cells treated with a 0.3 μM or 1 μM dose of 6-TG showed a ∼8.2% and ∼6.4% increase, respectively, of DNA in the tail compared to the untreated NUDT15 proficient cells and also 6-TG treated NUDT15 KO cells, although the total extent of DNA in the tail is lower at this time point compared to later time points (**Figure 7A**). After 48 h, the extent of strand breaks is statistically similar in NUDT15-proficient and deficient cells at both doses of 6-TG, although the comparisons are different by dose and across the time points examined. The NUDT15-proficient cells at the 0.3 μM dose showed a similar increase of ∼8.2% above non-treated parental cells, and this level persisted out to 72 h (**Figure 7B, C**). The NUDT15-proficient cells at the 1 μM dose showed a higher extent of strand breaks at 48 h, but a decrease ∼11.2% by 72 h, implying resolution of at least some of the strand breaks. In contrast, the extent of strand break induction in the NUDT15-deficient cells increased from 48 to 72 h at the 1 μM dose, an almost 4-fold increase from the same dose after 24 h of treatment (**Figure 7A, B**), p < 0.0001). The fold changes show that NUDT15-deficient cells accumulate a greater proportion of strand breaks that remain unresolved by 72 h compared to NUDT15-proficient cells (**Figure 7C**).

**Figure 7.**
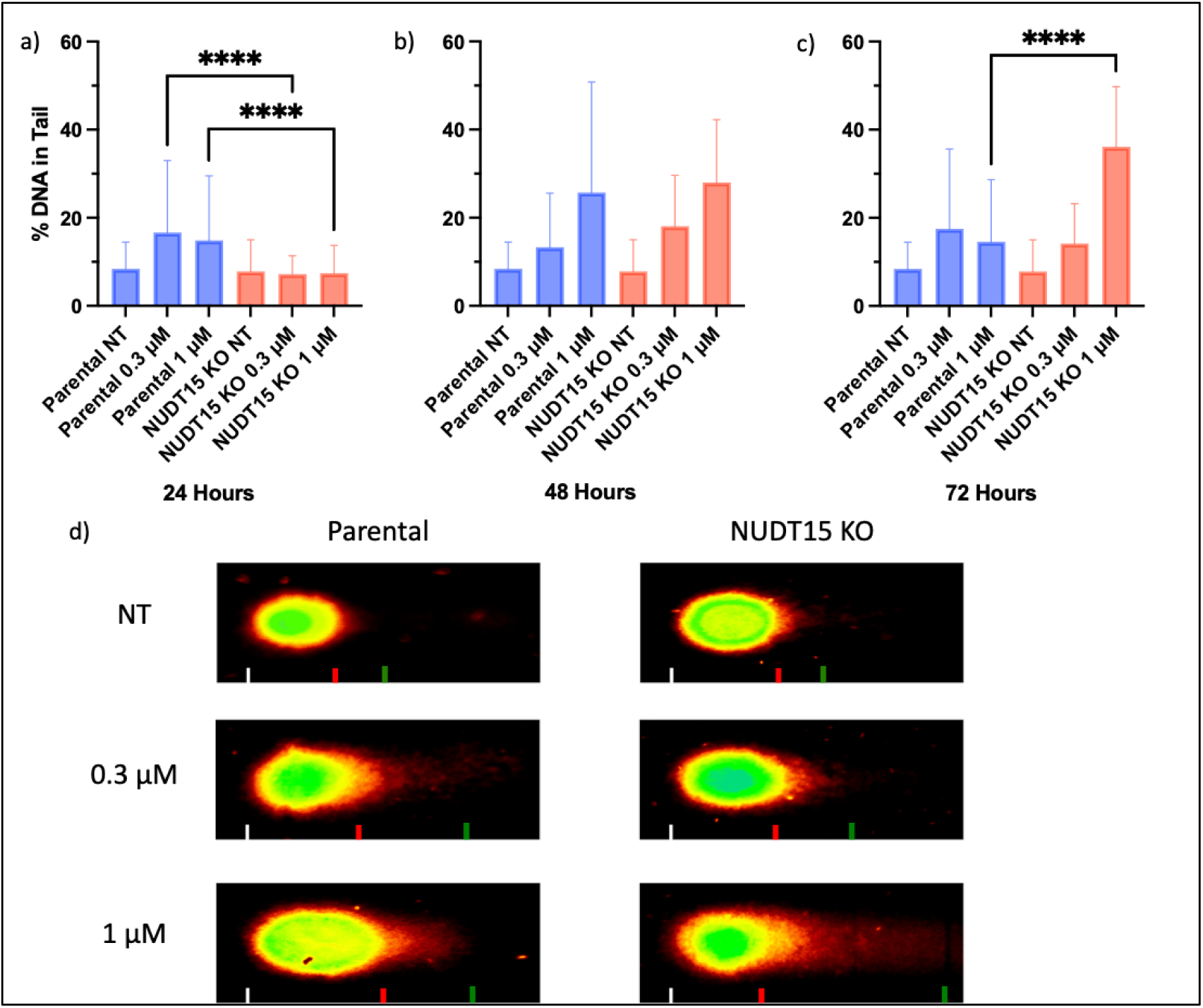
Comet assay measurement of DNA damage in 6-TG-treated parental and NUDT15 KO cells. The percent DNA in the tail for cells untreated (NT) or treated with 0.3 μM or 1 μM doses for **A)** 24 h, **B)** 48 h, and **C)** 72 h. **D)** Representative images of comets after 72 h NT, 0.3 μM, and 1 μM. Statistical significance was determined by two-way ANOVA **** p < 0.0001.

Next, chromosomal aberrations were measured following 6-TG treatments^24^. Aberrations scored included gaps/breaks, chromosome radials, fusions/end joining events, and complex exchanges, with examples shown in **Figure 8A**. NUDT15-proficient and deficient cells were treated with 0.3 μM or 1 μM of 6-TG for 6 h before recovering for 18 h and undergoing a 4 h colcemid incubation to enrich for mitotic cells. The NUDT15-proficient cells showed little change in chromosomal aberrations compared to the baseline of damage scored in the NT control (Figure 8B, bars 1-3). In contract, the NUDT15 KO cells showed increases in all four forms of scored aberrations with ∼5% and ∼8% of all counted chromosomes containing an aberration in the 0.3 μM and 1 μM 6-TG treated cells, respectively (**Figure 8B**, lanes 4-6). Note there was a striking increase in radials at the 1 μM dose in the NUDT15 KO cells (Figure 8B, lane 6, red). NUDT15 KO cells also show a complex exchange-type aberration in ∼1.6% and ∼2.1% of all scored chromosomes in the 6-TG treated cells. The formation of complex exchange aberrations is poorly understood but is presumed to be failed attempts at faithful repair and invocation of a secondary end-joining process that ligates broken fragments of chromatids or chromosomes (**Figure 8B**, lanes 5-6, purple). Slight increases in both gaps/breaks (differences of ∼1.6% and ∼0.7%) and fusions (differences of ∼1.0 % and ∼1.3%) were detected in the NUDT15 KO 6-TG treated cells compared to the NUDT15-proficient parental cells (**Figure 8B**, blue and green). The chromosomal aberration data supports the prior observations that a deficiency in NUDT15 contributes to chromosomal instability after treatments with 6-TG.

**Figure 8.**
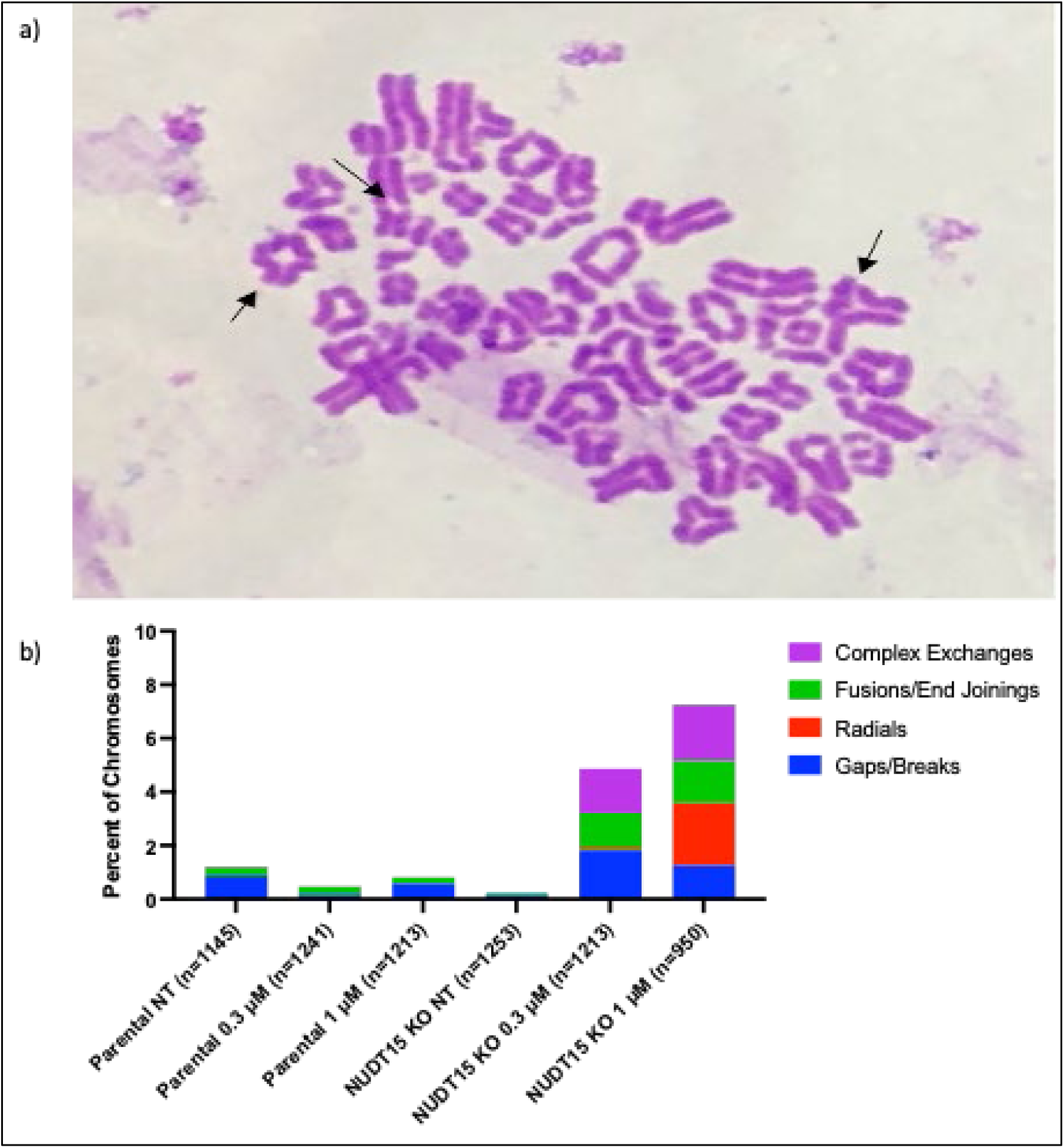
Chromosomal aberrations measured in parental and NUDT15 KO cells after 6-TG treatments. Metaphase spreads of parental and NUDT15 KO cells treated with 6-TG were generated and scored for chromosomal aberrations. **A)** Representative image of metaphase spreads with scorable aberrations. Arrows indicate examples of aberrations. From left to right arrows: Radial, Gap/Break, Complex Exchange. **B)** Quantification of aberrations in parental cells and NUDT15 KO cells after treatments with 0.3 μM or 1 μM 6-TG.

## Discussion

Thiopurines have been used as therapeutics for over 70 years and with indications beyond cancer to include use as immunosuppressants. The extensive clinical use of thiopurines has defined their therapeutic benefits and as well as risks of acute hematopoietic toxicity if the therapeutic window is exceeded and longer-term, less defined risk of iatrogenic cancers. Extensive clinical experience with thiopurines also uncovered the pharmacogenetic interactions of thiopurine-metabolizing enzymes and the biochemical and cellular basis of acute toxicity seen in those patients harboring homozygous germline mutations in those genes, including NUDT15^7–10^. In this study, we sought to explore a broader clinical cancer PGx utility, namely that NUDT15 deficiency represents a therapeutic vulnerability to target those cancer cells. We first confirmed that NUDT15 loss is a common co-occurrence with RB1-deleted cancers across many cancers, regardless of the frequency of RB1 loss in each cancer type. Interestingly, the association of NUDT15 and RB1 status also occurred in situations of gain particularly with a substantial subset of ovarian cancers, which we speculate would predict *resistance* to thiopurines.

We experimentally confirmed that NUDT15-defective cancer cells are sensitized to 6-TG, and that the mechanism of sensitization is consistent with its incorporation into DNA resulting in DNA damage and cell death. To better demonstrate the extent of the therapeutic window provided by NUDT15 deficiency in the setting of cyclical chemotherapy treatments used clinically, we used live cell imaging to detect changes in cell viability of a mixed culture of NUDT15-proficient and deficient cells over five passages and five cycles of 6-TG treatment (Figure 3). The imaging experiments demonstrate that NUDT15 loss causes a substantial therapeutic vulnerability to thiopurines compared to the NUDT15-proficient cells, which would include normal hematopoietic cells of a patient. While NUDT15 deficiency is known from clinical experience to cause hematopoietic toxicity^25^, and NUDT15-defective cancer cells were shown to be sensitized to 6-TG treatments^5^, the cytotoxic mechanism of action for 6-TG treatments in NUDT15-deficient cells remained unexplored. Because the established cytotoxic mechanism of action for thiopurines occurs through the induction of DNA damage^21, 23, 24, 26^, markers of DNA damage were measured and compared in the NUDT15-proficient and deficient cells. The induction of DNA damage is consistent with the known mechanism of action for thiopurines, further suggesting that the loss of NUDT15 exacerbates the DNA-directed effects of thiopurines. Of note, there was a striking difference in the level of 6-TG induced chromosomal aberrations, including dramatic increases in complex exchange events that were previously observed in HR-defective cells^23^. The work with cancer cells demonstrates that NUDT15-deficient cancer cells are sensitized to thiopurines and that the DNA-directed effects of thiopurines are exacerbated by NUDT15 deficiency. These observations further validate both the DNA-directed mechanism of action for thiopurines and the cellular activity of NUDT15, namely to inactivate the ultimate thiopurine metabolite, deoxy-6-thioGTP.

Importantly, we retrospectively analyzed thiopurine usage in males to examine if there might be an anti-cancer benefit associated their use, *i.e.*, whether clinical experience may support the hypothesis and pre-clinical results. Indeed, there is evidence suggesting that, among patients in the VA cohort, exposure to both thiopurines and allopurinol have a statistically significant reduced risk for prostate cancer development. Allopurinol usage was considered because of its known associations in affecting thiopurine metabolism. Indeed, it is appreciated clinically that allopurinol use can reduce hepatotoxicity from 6-MP in patients being treated for ALL^27^. Recent mass sequencing efforts of normal (preneoplastic) cells reveal that tissues not only harbor, but in many cases become colonized by, clones of cells carrying driver mutations in cancer genes long before the appearance of a fulminant detectable cancer^28^. We speculate that pre-cancer clones of RB1-deleted (and NUDT15 co-deleted) cells accumulate in the non- or pre-neoplastic cells of the prostate, and that thiopurine treatment (for whatever indication) wipes out these clones, eliminating them from being a platform on which new mutations and ultimately cancer develop. In other words, thiopurine usage for other indications serves as a fortuitous chemopreventive agent to select against prostate cells that genetically progress down the oncogenic pathway that loses RB1.

Understanding the pharmacogenetics of thiopurine metabolism and their mechanism of action in the context of pharmacogenetics serves to limit their cytotoxic effects in vulnerable patients with germline mutations in NUDT15. In this study, we propose that the same knowledge of thiopurine metabolism can be applied to personalized chemotherapy, in which tumors harboring somatically acquired mutations in drug metabolizing enzymes can be selectively targeted with those drugs that can exploit this vulnerability.

## Acknowledgment

The authors acknowledge Vitali Sikirzhytski, Ph.D. and the Microscopy Core of the COBRE Center for Targeted Therapeutics (CTT) for assistance with microscopy. Micah Goforth is acknowledged for assistance in scoring chromosomal abnormalities. The National Institutes of Health is acknowledged for funding (grant numbers by NIH COBRE grant (P20GM109091) for microscopy, R15 CA223956 (to M.D.W.), and R01DA054992 to S.S. and M.D.W. S.S. and J. M. are also supported by the South Carolina Center for Rural and Primary Healthcare for projects unrelated to this study. S.S. has received research grants from Boehringer Ingelheim, Coherus BioSciences, EMD Serono, and Alexion Pharmaceuticals, all for projects unrelated to study.

## Author contributions

J. M. performed all experiments and data analysis with cancer cells. J. M. performed the analysis of VA VINCI data, with oversight from S.S. P.B. performed cancer genome analysis. M.W. conceived of the idea, supervised the research, helped design experiments, and analyzed the data. All authors contributed to writing and editing the manuscript and approved the final draft of the manuscript.

